# B7-H3 upregulation in ischemic stroke: friend or foe?

**DOI:** 10.64898/2025.11.28.691244

**Authors:** Siva Reddy Challa, Isidra M. Baker, Casimir A. Fornal, Sahil Reddy Mada, Nabeeha Khan, Samantha Jackson, Erick Saldes, Jeffrey D. Klopfenstein, Swapna Asuthkar, Krishna Kumar Veeravalli

**Author notes:** Corresponding author at: Department of Cancer Biology and Pharmacology, University of Illinois College of Medicine Peoria, 1 Illini Drive, Peoria, IL 61605, USA. E-mail address (K.K. Veeravalli). Shared first authorship (Siva Reddy Challa, Isidra M. Baker).

## Abstract

B7-H3 (CD276) is an immune checkpoint co-signaling molecule expressed on immune and non-immune cells. It is best known for suppressing T-cell responses but can also promote inflammation depending on the microenvironment. In neuroinflammatory models such as experimental autoimmune encephalomyelitis, B7H3 expression increases concomitantly with the inflammatory response, and its inhibition is associated with reduced disease progression. Although its role in ischemic stroke remains unclear, we hypothesized that cerebral ischemia/reperfusion (I/R) would upregulate B7-H3 expression in the ischemic brain and that increased B7-H3 expression would positively correlate with pro-inflammatory cytokine expression. Young and aged male and female rodents, including normotensive and spontaneously hypertensive rats to model comorbid hypertension, underwent transient middle cerebral artery occlusion (MCAO) followed by reperfusion. Brain tissue was collected on post-ischemic days 1, 3, 5, or 7. B7-H3 mRNA was analyzed by real-time PCR, whereas protein expression was assessed by Western blotting and immunohistochemistry at selected time points. B7-H3 expression was significantly upregulated in the ischemic brain across sexes, age groups, and species. The extent of B7-H3 degradation in the ischemic brain was influenced by species, sex, age, and time after cerebral I/R. Upregulation of B7-H3 was observed at both the mRNA and protein levels, and increased expression was localized primarily to the somatosensory cortex and caudate putamen in the ipsilateral hemisphere, the main regions affected in this MCAO model. Elevated B7-H3 expression in the ischemic brain positively correlated with the pro-inflammatory mediator TNFα. The temporal profile of B7-H3 expression observed in rats paralleled the early inflammatory phase associated with secondary tissue damage following ischemic stroke. These findings identify B7-H3 as an ischemia-induced immune checkpoint molecule in the brain that may modulate post-stroke immune responses and support further investigation into its beneficial versus detrimental roles in neuroinflammation, as well as its potential as a therapeutic target following cerebral I/R.

## 1. Introduction

Stroke is a leading cause of long-term neurological deficits and disability worldwide. It is currently the fifth leading cause of death in the United States and the second leading cause of death globally (Furie, 2020). Ischemic stroke remains the most prevalent type, accounting for approximately 87% of all strokes (Virani et al., 2020). The only FDA-approved pharmacological therapies for a subset of patients with acute ischemic stroke (AIS) are thrombolytic agents that remove intravascular clots, namely tissue-type plasminogen activator (tPA) and tenecteplase (TNK). There are no FDA-approved treatments for AIS patients beyond clot removal. In addition, drugs that mitigate progressive brain injury or enhance functional recovery following recanalization by thrombolysis or endovascular thrombectomy remain an unmet need. Given the global burden of stroke, it is clinically important to identify novel targets and develop therapeutic strategies that improve outcomes in AIS patients.

Advanced age and vascular comorbidities such as hypertension are major risk factors that increase both the incidence and severity of ischemic stroke (Fisher et al., 2007; Howells et al., 2010; O’Donnell et al., 2010; Mozaffarian et al., 2015). Hypertensive animals and aged rodents therefore provide clinically relevant models for investigating mechanisms that differentially modulate stroke injury and recovery in high-risk populations.

Harmful neuroinflammation triggered by aberrant immune activation following cerebral ischemia/reperfusion (I/R) critically exacerbates brain damage and impedes functional recovery (Jayaraj et al., 2019). Analysis of the GEO database indicates that several immune-related genes, including B7-H3, are highly expressed in stroke patients (Tao et al., 2022). B7-H3 (CD276) is an immune checkpoint co-signaling molecule expressed on both immune and non-immune cells that plays a multifaceted role at the interface of innate and adaptive immunity. B7-H3 can function as both a T-cell costimulator and coinhibitor. As a costimulatory molecule, B7-H3 enhances T-cell proliferation and IFNγ production and augments pro-inflammatory cytokine release from monocytes/macrophages (Chapoval et al., 2001; Zhang et al., 2010). Recent research shows that upregulation of B7-H3 augments inflammatory signaling pathways, resulting in elevated cytokine production and breakdown of the blood-brain barrier (Chen et al., 2017). Consistent with these findings, B7-H3 knockout mice exhibit reduced inflammation in models of collagen-induced arthritis and experimental autoimmune encephalomyelitis (Luo et al., 2015). In contrast, B7-H3 functions predominantly as a coinhibitory molecule in the context of cancer (Castellanos et al., 2017; Kontos et al., 2021; Getu et al., 2023). These reports clearly illustrate the functional duality of B7-H3. It may exert either costimulatory or coinhibitory effects in different environments by differentially modulating distinct T-cell subsets. For example, B7-H3 enhances Th1/Th17 responses and suppresses Th2 responses, depending on the predominant T-cell subsets present in the microenvironment (Luo et al., 2015). Despite the known role of B7-H3 in immune regulation and its detection in stroke patient samples, no studies have systematically characterized B7-H3 expression in preclinical stroke models across different rodent species, sexes, ages, and comorbidities, and neither its temporal expression profile nor its regional localization has been determined.

This study aims to characterize B7-H3 expression in rodent stroke models across species, sex, age, and post-ischemic time points following cerebral I/R. By defining the temporal expression profiles and regional localization of B7-H3 upregulation in these models, we aim to identify suitable preclinical rodent models and to delineate an optimal treatment window for future evaluation of therapies targeting B7-H3. We also examined B7-H3 expression in normotensive Wistar-Kyoto (WKY) rats and spontaneously hypertensive rats (SHRs) to assess whether B7-H3 upregulation is accentuated in hypertensive animals that develop more severe neurological deficits after middle cerebral artery occlusion (MCAO). In addition, we analyzed the relationship between B7-H3 expression and the expression levels of three pro-inflammatory cytokines (IL-1β, IL-6, and TNFα) in the ischemic brain. We found that B7-H3 expression was upregulated in the ipsilateral (ischemic) hemisphere in both rats and mice, irrespective of age or sex, and was further accentuated in hypertensive rats compared with normotensive controls. In young Sprague-Dawley rats, B7-H3 expression in the ipsilateral (ischemic) hemisphere remained elevated for at least 5 days after cerebral I/R. Although all three predominant B7-H3 isoforms (57, 53, and 34 kDa) were consistently detected in the ischemic brains of rats and mice, the relative degradation status of B7-H3, as reflected by the 57/34 kDa isoform ratio, was modulated by species, sex, age, and time after ischemia. Moreover, B7-H3 expression showed a significant positive correlation with TNFα expression, consistent with a potential link between B7-H3 and post-ischemic neuroinflammation.

## 2. Material and methods

### 2.1. Ethics and compliance statements

All animal procedures were conducted in accordance with a protocol approved by the Institutional Animal Care and Use Committee (IACUC) at the University of Illinois College of Medicine Peoria (UICOMP). All experiments adhered to the scientific, humane, and ethical principles of UICOMP and to the guidelines outlined in the *Guide for the Care and Use of Laboratory Animals* (NIH Publication No. 86-23, revised; U.S. Department of Health and Human Services). Animal experiments were designed, conducted, and reported in accordance with the Animal Research: Reporting of In Vivo Experiments (ARRIVE) guidelines (Percie du Sert et al., 2020).

### 2.2. Animals and experimental groups

Sprague-Dawley (SD) rats were obtained from Envigo (Indianapolis, IN, USA). Wistar-Kyoto (WKY) rats and spontaneously hypertensive rats (SHRs) were obtained from Charles River (Wilmington, MA, USA). C57BL/6J mice were obtained from The Jackson Laboratory (Bar Harbor, ME, USA). Both rats and mice were housed in the UICOMP Laboratory Animal Care Facility and maintained under controlled temperature and humidity on a 12-h light-dark cycle with *ad libitum* access to food and water. A total of 59 young (3-month-old) male SD rats, 14 young (3-month-old) male WKY rats, 24 young (3-month-old) male SHRs, and 59 C57BL/6J mice (both sexes combined; young [3-month-old] and aged [17-19-month-old]) were used in this study and were randomly assigned to either Control or Stroke groups, as summarized in **Table 1**.

**Table 1.**
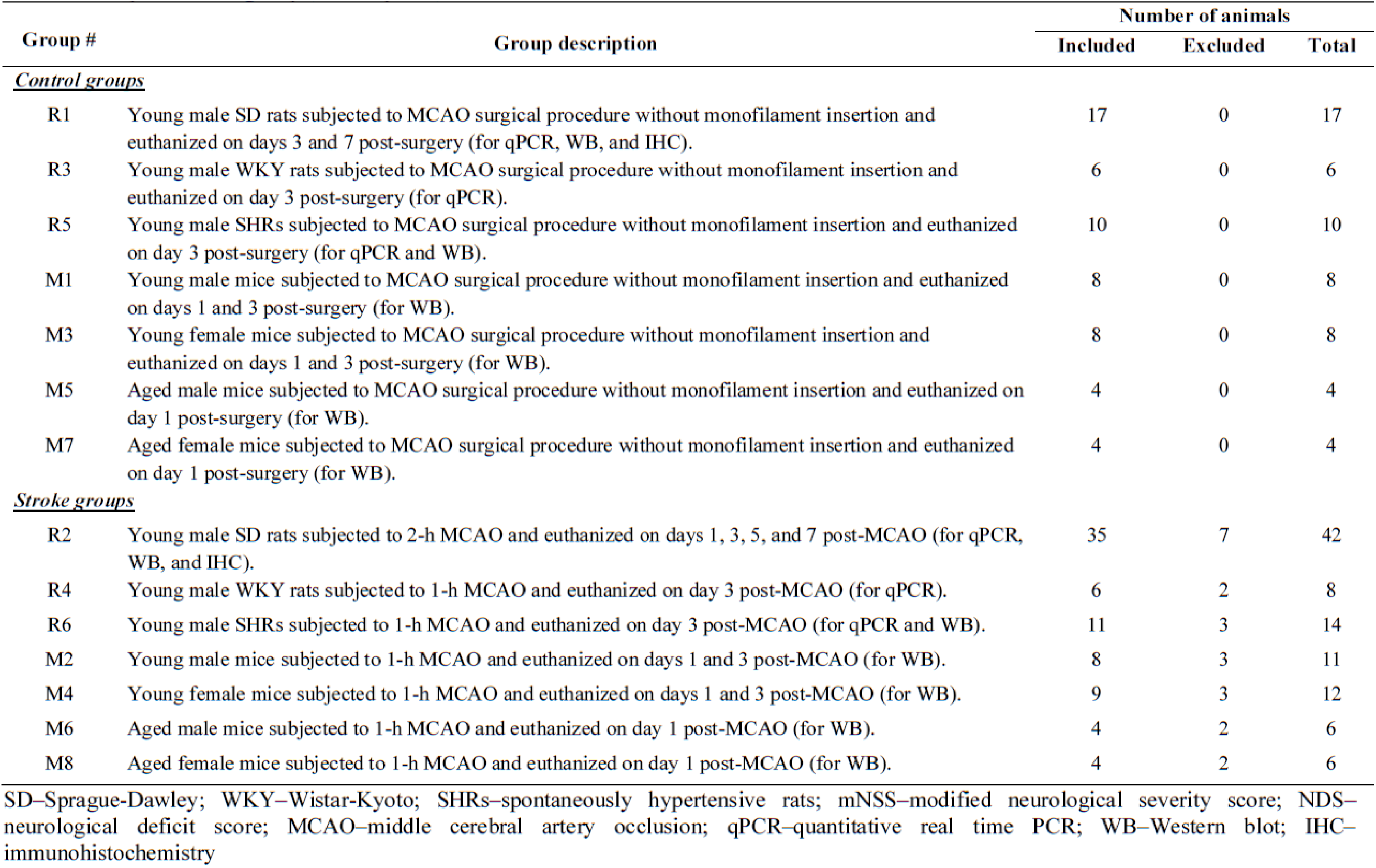
Experimental groups, description, and animal numbers.

### 2.3. Stroke induction in rats

Stroke induction in rats was performed by right transient middle cerebral artery occlusion (MCAO) using the intraluminal monofilament suture technique. Rats were deeply anaesthetized with 3.0-3.5% isoflurane delivered via a VetFlo isoflurane anesthesia system (Kent Scientific Corporation, Torrington, CT, USA) and maintained at 2.5-3.0% isoflurane throughout the surgery. Under anesthesia, rats were placed on a SurgiSuite multifunctional surgical platform with integrated far-infrared warming (Kent Scientific Corporation, Torrington, CT, USA) to maintain normothermia and prevent hypothermia during surgery. An aseptic technique was followed throughout the procedure. A ventral midline neck incision (∼25 mm) was made, and the right common carotid artery (CCA), internal carotid artery (ICA), and external carotid artery (ECA) were surgically exposed. The ECA was permanently ligated rostrally with a single 5-0 silk suture, and a second loose ligature was placed around the ECA near the CCA bifurcation. Microaneurysm clips were applied to the CCA and ICA. The ECA was then transected between the two ligatures. An appropriately sized silicone rubber-coated monofilament suture (Doccol Corporation, Sharon, MA, USA) was introduced through the stump of the severed ECA and advanced into the ICA. The microaneurysm clip was removed from the ICA, and the monofilament was gently advanced 19-20 mm to a premarked distance corresponding to the origin of the middle cerebral artery (MCA). The loose ligature at the bifurcation was then tightened around the ECA containing the monofilament. The microaneurysm clip on the CCA was removed, and the neck incision was closed with surgical wound clips. To restore blood flow, reperfusion was initiated 2 hours after MCAO in SD rats and 1 hour after MCAO in WKY rats and SHRs. The neck incision was reopened by removing the wound clips, and a microaneurysm clip was reapplied to the CCA. The ECA ligature was loosened, the monofilament suture was carefully withdrawn, and the ligature was retied to achieve hemostasis. The microaneurysm clip on the CCA was then removed, and the neck incision was closed with 3-0 nylon sutures. Rats in the control group underwent surgical procedures identical to the MCAO surgery, except that the monofilament was not inserted. Throughout this manuscript, “control” refers to these sham-operated animals.

Pre- and post-surgical care was provided in accordance with the IMPROVE guidelines (*Ischemia Models: Procedural Refinements of In Vivo Experiments*) (Percie du Sert et al., 2017). Post-surgical care for rats included administration of the analgesic carprofen (5 mg/kg, s.c., once daily for 2 days following surgery), and subcutaneous sterile saline (approximately 2 mL on the day of surgery and for 2 days thereafter) to maintain hydration. A topical triple-antibiotic ointment containing bacitracin, neomycin, and polymyxin B was applied prophylactically to the neck incision to prevent infection. In addition, moistened chow was placed on the cage floor to facilitate access to food during recovery.

### 2.4. Modified Neurological Severity Score (mNSS) assessment in rats

The mNSS is a composite index of motor, sensory, reflex and balance function (Chen et al., 2001). Neurological deficits were assessed using the mNSS in rats at 2-4 h and again on post-MCAO day 1 to determine the severity of stroke injury. Scores range from 0 (no deficit) to 18 (maximal neurological impairment).

### 2.5. Stroke induction in mice

Stroke induction in mice was performed using the same right transient MCAO monofilament technique as in rats, with minor modifications. Mice were deeply anaesthetized with 3% isoflurane delivered via a VetFlo isoflurane anesthesia system (Kent Scientific Corporation, Torrington, CT, USA) and maintained at 2% isoflurane throughout the surgery. Under anesthesia, mice were placed on a SurgiSuite multifunctional surgical platform with integrated far-infrared warming (Kent Scientific Corporation, Torrington, CT, USA) to maintain normothermia and prevent hypothermia during surgery. An aseptic technique was followed throughout the procedure. A ventral midline neck incision (10-15 mm) was made, and the right CCA, ICA, and ECA were surgically exposed. The ECA was permanently ligated rostrally with a single 6-0 silk suture, and a second loose ligature was placed around the ECA near the CCA bifurcation. Blood flow in the CCA and ICA was temporarily occluded by lifting each vessel with 5-0 silk sutures. The ECA was then transected between the two ligatures. An appropriately sized silicone rubber-coated monofilament suture (Doccol Corporation, Sharon, MA, USA) was introduced through the stump of the severed ECA and advanced into the ICA while tension on the ICA ligature was slowly released. The monofilament was gently advanced approximately 9 mm to a premarked distance corresponding to the origin of MCA. The loose ligature was then tightened around the ECA stump containing the monofilament. Tension on the CCA was relieved by removing the silk suture, and the skin incision was temporarily closed with surgical wound clips. Following completion of the initial surgery, mice were allowed to recover from anesthesia in a surgical recovery cage equipped with a water-perfused heating pad beneath one half of the cage to allow behavioral thermoregulation. Mice were re-anesthetized with isoflurane shortly before the designated reperfusion time (1 hour after MCAO). The neck incision was reopened by removing the wound clips, and the CCA was again lifted with a 5-0 silk suture to temporarily block blood flow. The ECA ligature was loosened, the monofilament suture was carefully withdrawn to initiate reperfusion, and the ligature was retightened to achieve hemostasis. Tension on the CCA was then relieved and the neck incision was closed with 5-0 nylon sutures. Mice in the control group underwent surgical procedures identical to the MCAO surgery, except that the monofilament was not inserted. Thus, “control” mice are refer to sham-operated animals.

Pre- and post-surgical care was provided in accordance with the IMPROVE guidelines (*Ischemia Models: Procedural Refinements of In Vivo Experiments*) (Percie du Sert et al., 2017). Post-surgical care for mice included administration of the analgesic carprofen (5 mg/kg, s.c., once daily for 2 days following surgery), and subcutaneous sterile saline (approximately 0.5 mL on the day of surgery and for 2 days thereafter) to maintain hydration. A topical triple-antibiotic ointment containing bacitracin, neomycin, and polymyxin B was applied prophylactically to the neck incision to prevent infection. In addition, moistened chow was placed on the cage floor to facilitate access to food during recovery.

### 2.6. Neurological Deficit Score (NDS) assessment in mice

Neurological deficit score (NDS) assessments were conducted in mice at 2-4 h and again on post-MCAO day 1 to determine the severity of stroke injury. Neurological function was graded on a 0-3 scale as follows: 0—no observable deficits; 1—forelimb flexion; 2—forelimb flexion and decreased resistance to lateral push; 3—forelimb flexion, decreased resistance to lateral push, and circling.

### 2.7. Exclusion criteria

Animals that died or were euthanized during the study period, as well as those exhibiting post-mortem bleeding in the region of the MCA, were excluded from the study. In addition, mice with NDS scores < 2 at either 2-4 hours or post-MCAO day 1 were excluded. Finally, any data points identified as outliers by Grubb’s test (α = 0.01) were excluded from the analysis.

### 2.8. Tissue collection and processing

Rats from both control and stroke groups were used for tissue collection on days 1, 3, 5, or 7 after MCAO, and mice on days 1 and 3 after MCAO. Under deep anesthesia, animals were transcardially perfused, and brains were collected for molecular and histological analyses as described below.

For real-time PCR and Western blot analysis in rats, animals were perfused intracardially with ice-cold phosphate-buffered saline (PBS; 1x) to remove intravascular blood. Brains were rapidly removed, and the ipsilateral hemispheres were separated from the contralateral hemispheric tissue and stored at -80 °C until processing for RNA isolation and protein extraction. For immunohistochemical analysis in rats, animals were perfused intracardially with PBS followed by 10% neutral buffered formalin. Brains were post-fixed in the same fixative for 24 hours at 4 °C and then cryoprotected in 30% (w/v) sucrose in PBS until they sank. Cryoprotected brains were embedded in optimal cutting temperature (OCT) compound (Tissue-Tek OCT, Catalog # 4583; Sakura Finetek USA, Torrance, CA, USA), frozen, and stored at -80 °C until sectioning. Coronal brain sections (40 µm thick) were cut on a cryostat (Leica CM1950; Leica Biosystems), and sections containing the caudate-putamen at the level of the anterior commissure were collected into 24-well polystyrene plates containing PBS with 0.02% sodium azide (as a preservative) and stored at 4 °C for immunohistochemical staining with DAB.

For Western blot analysis in mice, animals were perfused intracardially with ice-cold PBS in the same manner. Brains were removed, the ipsilateral hemisphere was dissected, and tissue was stored at -80 °C until protein extraction for Western blot analysis.

### 2.9. RNA isolation and cDNA synthesis

Total RNA was extracted from the entire ipsilateral cerebral hemispheres of rats from both control and stroke groups using TRIzol Reagent (Invitrogen, Carlsbad, USA). One microgram of total RNA from each sample was reverse transcribed into cDNA using iScript cDNA Synthesis Kit (Bio-Rad Laboratories, Hercules, CA, USA), and the resulting cDNA was diluted 1:10 and stored at -20 °C for subsequent analysis.

### 2.10. Real-time PCR analysis

For each diluted (1:10) cDNA sample, reactions were assembled using the iTaq Universal SYBR Green Supermix (Bio-Rad Laboratories, Hercules, CA, USA) according to the manufacturer’s instructions. Forward and reverse primers for the target genes (Integrated DNA Technologies, Coralville, IA, USA) were diluted 1:10 in nuclease-free water. Primer sequences for the target genes are listed in **Table 2**. PCR reactions were performed in triplicates using the following thermal cycling conditions: initial denaturation at 95°C for 5 min; 40 cycles of 95 °C for 30 sec, 60 °C for 30 sec, and 72 °C for 30 sec; and a final extension at 72 °C for 5 min. Reactions were run on an iCycler IQ Multi-Color real-time PCR detection system (Bio-Rad Laboratories, Hercules, CA, USA). Data was collected and recorded using iCycler IQ software (Bio-Rad Laboratories, Hercules, CA, USA) and expressed as threshold cycle (Ct) values, defined as the number of cycles at which the fluorescent intensity of the SYBR Green dye is significantly above background fluorescence. For each sample, the mean Ct values of the triplicates, after removal of any outlier, was used for analysis. *18S* rRNA served as the internal reference gene. Relative quantification of target gene expression was normalized to *18S* rRNA, and fold change in target gene expression in test samples relative to control samples was calculated using the 2^ΔCt method, where fold change = 2^(ΔCt control)/2^(ΔCt test).

**Table 2.**
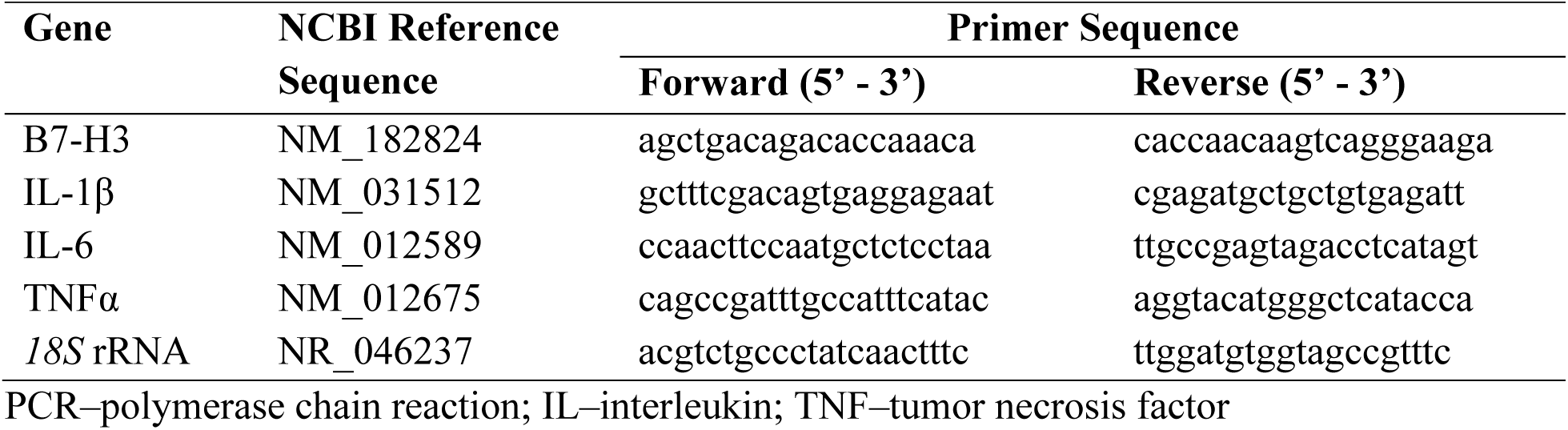
Rat primers used for real-time PCR analysis.

### 2.11. Western blot analysis

Westen blot analysis was performed using brain tissue lysates obtained on post-MCAO days 1 or 3 from rats and mice in both control and stroke groups. Equal amounts of protein from each sample were subjected to SDS-PAGE and immunoblotting with a mouse monoclonal anti-B7-H3 antibody (Catalog # sc-376769; Santa Cruz Biotechnology, USA) followed by an HRP-conjugated goat anti-mouse IgG secondary antibody. Each immunoblot was then reprobed with a mouse monoclonal anti-GAPDH antibody (Catalog # sc-32233; Santa Cruz Biotechnology, USA) followed by incubation with the same HRP-conjugated goat anti-mouse IgG secondary antibody. Immunoreactive bands were visualized using an enhanced chemiluminescence (ECL) Western blot detection reagent (Bio-Rad Laboratories, USA). Band intensities were quantified using NIH ImageJ software (version 1.54p) and normalized to the loading control, GAPDH.

### 2.12. Immunohistochemistry

For each rat, a single coronal brain section through the caudate-putamen at the level of the anterior commissure was processed for immunohistochemistry. Coronal sections were mounted onto adhesive microscope slides (Catalog #1354W; Diamond White Glass, Globe Scientific, Mahwah, NJ, USA) from 50 mM phosphate buffer (pH 7.4). Slides were dried overnight at 37°C and then left at room temperature for an additional day prior to processing. Slide-mounted sections were subjected to heat-induced epitope retrieval in 10 mM citric acid (pH 6.0) at 90°C for 30 min using a Precision water bath (Model 282; Winchester, VA, USA). Following rinses with PBS (1×), sections were incubated in 0.3% hydrogen peroxide in PBS for 30 min to quench endogenous peroxidase activity. Sections were then permeabilized with 0.1% Tween-20 in PBS and blocked with 5% normal goat serum containing 0.05% Triton X-100 in PBS. Following blocking, sections were incubated overnight at 4°C with a mouse monoclonal anti-B7-H3 antibody (Catalog # sc-376769; Santa Cruz Biotechnology, USA) at a 1:100 dilution in 3% normal goat serum containing 0.05% Tween-20 in PBS. The following day, sections were rinsed with PBS and incubated for 1 h with an HRP polymer-conjugated goat anti-mouse IgG secondary antibody (Catalog #VC001; R&D Systems, Minneapolis, MN, USA). After additional rinses with PBS, sections were reacted with 3,3′-diaminobenzidine (DAB) using the SigmaFAST tablet set (Catalog #D4418; Sigma-Aldrich, St. Louis, MO, USA) containing urea hydrogen peroxide. Once optimal DAB staining intensity was achieved, the reaction was stopped by rinsing the sections in deionized water. Slides were dehydrated through graded ethanol series (50%, 70%, 95%, and 100%), cleared in two changes of xylene, and coverslipped with DPX mounting medium (Catalog #360294H; VWR, USA).

After the slides were air-dried, they were cleaned and scanned at 10,000 dpi using a PrimeHisto XE slide scanner (Pacific Image Electronics, Taiwan). Digital images were analyzed using NIH ImageJ software (version 1.54p). Each image was imported and converted to 8-bit format. The integrated density of B7-H3 staining (stained area x mean grayscale value) was quantified separately in the contralateral (non-ischemic) and ipsilateral (ischemic) hemispheres. A fixed threshold range (minimum and maximum grayscale values) corresponding to the B7-H3-positive signal was established and applied uniformly to all sections to ensure consistency and comparability between treatment groups. Because B-H3-positive staining appeared as darker pixels with lower grayscale values, the measured grayscale values within the threshold region were numerically inverted so that higher values corresponded to stronger staining. These inverted values were used for data analysis.

### 2.13. Statistical analysis

Statistical analysis of the data was performed using *GraphPad Prism 10.4.2* for Windows (GraphPad Software, San Diego, CA, USA). Outliers in the data were identified using Grubb’s test (α = 0.01) and excluded from the analysis. Quantitative data from each experiment were tested for normality (Shapiro-Wilk test) and equality of variances (F-test and Bartlett’s test). Based on the number of groups present in each experiment and the outcome of the normality and variance tests, appropriate statistical tests were applied, including Mann-Whitney test when normality was not met, two-tailed unpaired t-test with Welch’s correction when variances were unequal, one-way ANOVA followed by Dunnett’s multiple comparisons test, and two-way ANOVA followed by Sidak’s multiple comparisons test. Pearson’s correlation test (Pearson’s *r*) was used to assess relationships between variables. Differences between groups were considered statistically significant at *p* < 0.05. All data are expressed as mean ± SEM.

## 3. Results

### 3.1. B7-H3 expression increases in the brain after ischemic stroke in young rats

Cerebral ischemia/reperfusion (I/R) induced by transient middle cerebral artery occlusion (MCAO) in young male Sprague-Dawley (SD) rats resulted in a time-dependent increase in B7-H3 mRNA expression in the ipsilateral brain with levels higher on post-stroke days 1, 3, 5, and 7 compared with the control group (**Fig. 1A**). B7-H3 upregulation was modest on day 1, peaked on day 3, declined but remained elevated on day 5, and further declined toward baseline by day 7. However, one-way ANOVA followed by Dunnett’s multiple comparisons test showed that these increases were statistically significant only on day 3 and 5 (both *p* < 0.0001), whereas changes on days 1 and 7 did not reach significance. Western blot analysis of ipsilateral brain samples collected on day 3 post-MCAO demonstrated increased protein expression of all B7-H3 isoforms in the stroke group relative to controls (**Fig. 1B**). A two-tailed unpaired t-test confirmed that the 57-kDa isoform was significantly increased (*p* = 0.0363) and the 53-kDa and 34-kDa isoforms were markedly upregulated (both *p* < 0.0001) in stroke brains compared with the control group.

**Fig. 1.**
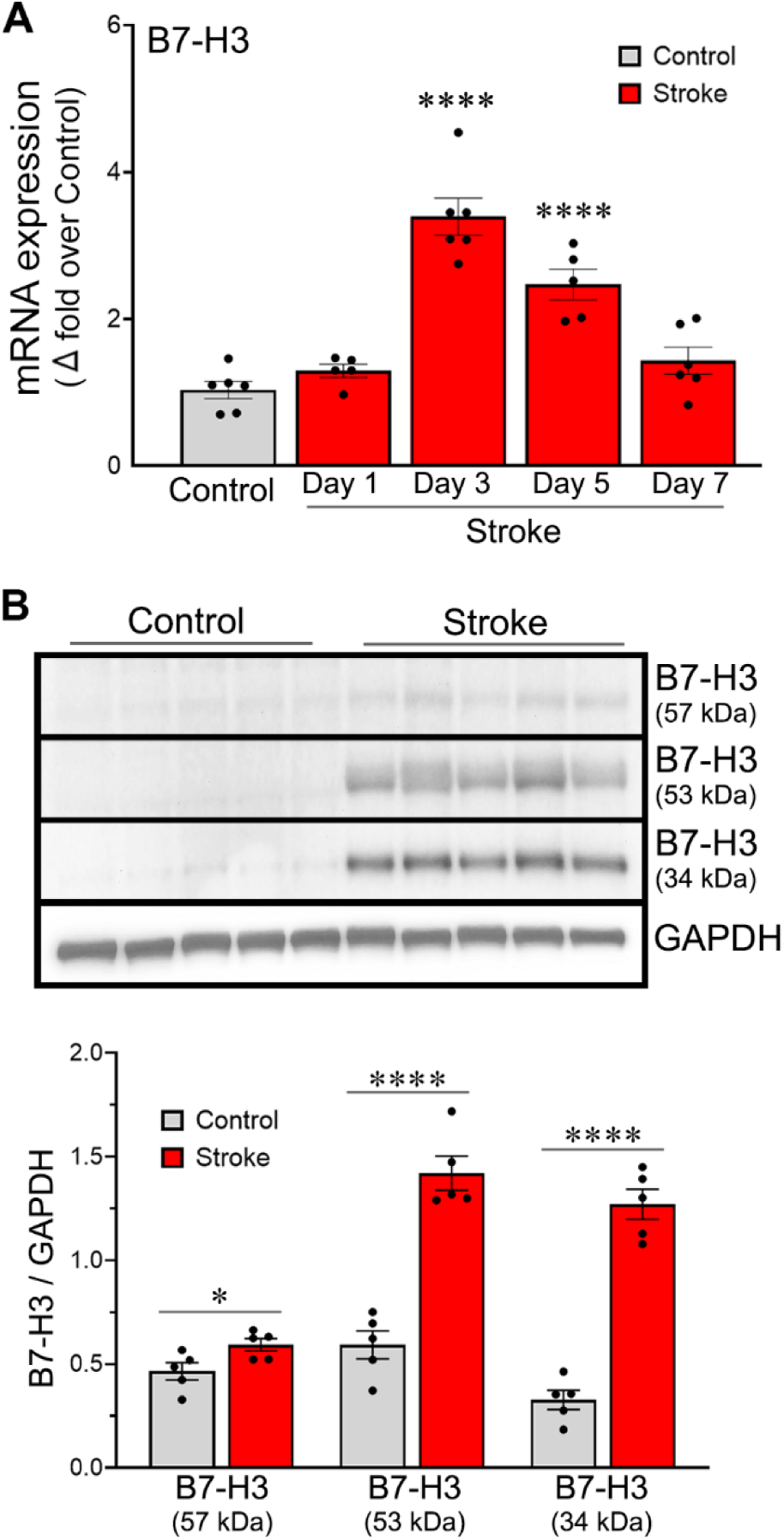
B7-H3 expression increases in the brain following ischemic stroke in young rats. (A) Column scatter plot shows B7-H3 mRNA expression in the ipsilateral brain on post-ischemic days 1, 3, 5, and 7. Young rats were subjected to 2-hour MCAO followed by reperfusion or sham surgery, and PBS-perfused brain tissue was collected at the indicated time points. Error bars indicate SEM; n=5–6/group. \*\*\*\**p*<0.0001 versus control. (B) Immunoblot analysis demonstrates upregulation of three B7-H3 isoforms (57, 53, and 34 kDa) in the ischemic brains of rats on day 3 post-MCAO. Column scatter plot shows the quantified expression of the 57-, 53-, and 34-kDa B7-H3 isoforms relative to GAPDH expression in PBS-perfused ipsilateral brain tissue. Error bars indicate SEM; n=5/group. \**p*<0.05, \*\*\*\**p*<0.0001.

### 3.2. B7-H3 expression increases in the ischemic brain of both sexes of young mice

To determine whether the ischemia-induced B7-H3 upregulation observed in rats extends to a second rodent species, we examined B7-H3 protein expression in the ipsilateral brains of young male and female mice subjected to 1-hour MCAO. Western blot analysis of brain samples collected on post-stroke days 1 and 3 showed increased expression of all B7-H3 isoforms in the stroke groups compared with their respective control groups (**Fig. 2**). In young males, a two-tailed unpaired t-test (with Welch’s correction applied when variances were unequal) revealed significantly higher B7-H3 protein levels in the stroke group on both days 1 and 3 compared with controls (day 1: *p* = 0.0009 for the 57-kDa isoform, *p* < 0.0001 for the 53-kDa isoform, and *p* = 0.0008 for the 34-kDa isoform; day 3: *p* < 0.0001 for all isoforms) (**Fig. 2A**). Similarly, in young females, a two-tailed unpaired t-test (with Welch’s correction applied when variances were unequal) showed significant increases in B7-H3 protein expression in stroke brains compared with controls (day 1: *p* = 0.0046 for the 57-kDa isoform and *p* = 0.0024 for the 34-kDa isoform; day 3: *p* = 0.0005 for the 57-kDa isoform, *p* = 0.0002 for the 53-kDa isoform, and *p* < 0.0001 for the 34-kDa isoform), whereas the 53-kDa isoform on day 1 did not reach statistical significance (*p* = 0.0504) (**Fig. 2B**).

**Fig. 2.**
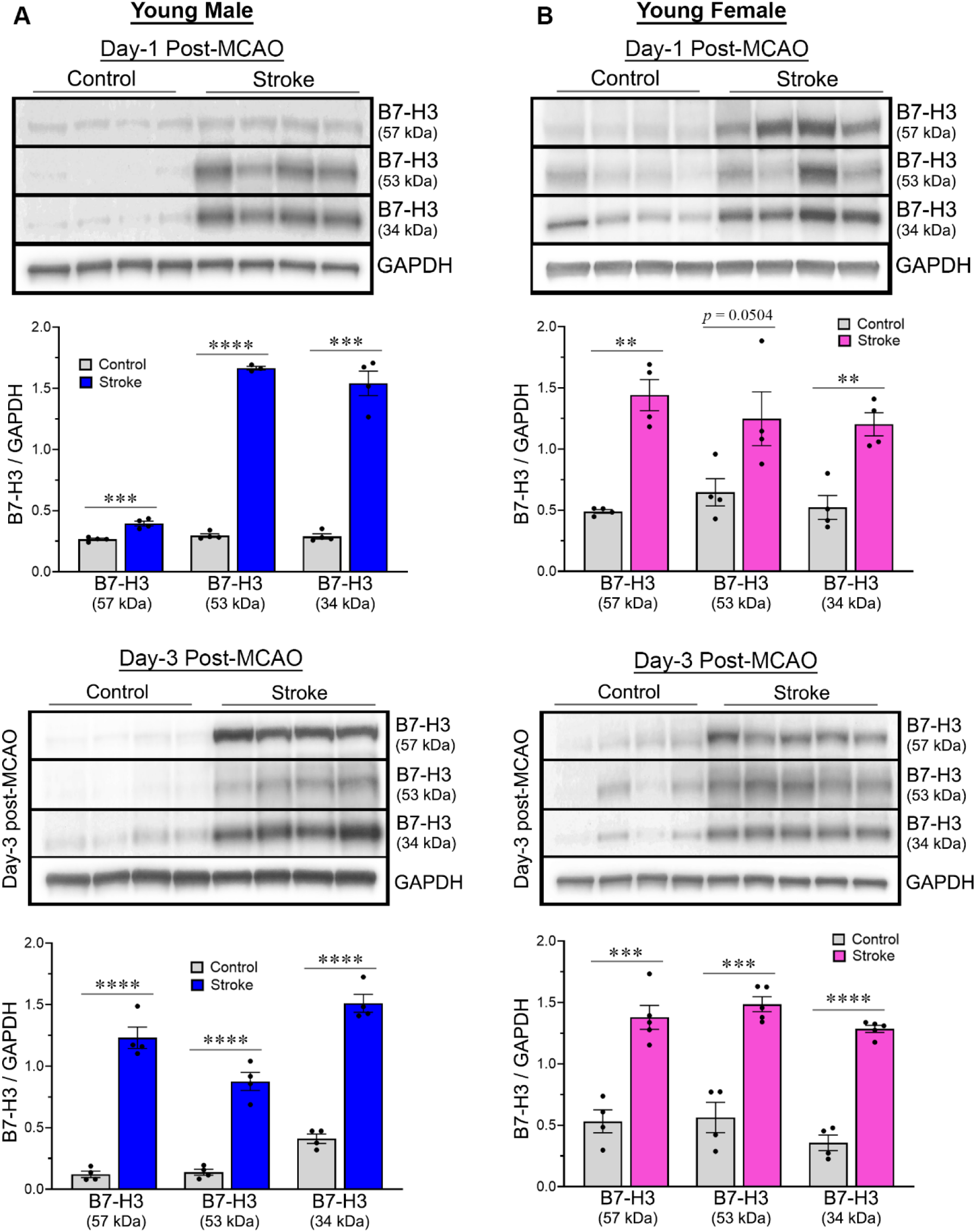
B7-H3 expression in the brain after ischemic stroke in young male and female mice. Immunoblots demonstrate protein expression of three B7-H3 isoforms (57, 53, and 34 kDa) in the ipsilateral (ischemic) brains of young male (A) and female (B) mice compared with age-matched controls. Both sexes were subjected to 1-hour MCAO followed by reperfusion, and PBS-perfused brain tissue was collected on post-ischemic days 1 and 3. Column scatter plots show the quantified expression of the 57-, 53-, and 34-kDa B7-H3 isoforms relative to GAPDH expression. Error bars indicate SEM; n=4–5/group. \*\**p*<0.01, \*\*\**p*<0.001, \*\*\*\**p*<0.0001.

### 3.3. Upregulation of B7-H3 expression in the ischemic brains of hypertensive and aged rodent models

Because advanced age, hypertension, and biological sex all influence stroke risk and severity, we next examined B7-H3 protein expression in more clinically relevant models: spontaneously hypertensive rats (SHRs) and aged male and female mice subjected to 1-hour MCAO. Western blot analysis of ipsilateral brain samples collected on post-stroke day 1 from aged male and female mice showed expression of all B7-H3 isoforms in the stroke groups compared with their respective control groups (**Fig. 3A, B**). In aged males, two-tailed unpaired t-tests (with Welch’s correction applied when variances were unequal) revealed significantly higher B7-H3 protein levels in the stroke group than in controls (*p* = 0.0006 for the 57-kDa isoform, *p* = 0.0037 for the 53-kDa isoform, and *p* < 0.0001 for the 34-kDa isoform) (**Fig. 3C**). Similarly, in aged females, two-tailed unpaired t-tests (with Welch’s correction applied when variances were unequal) demonstrated significant increases in B7-H3 protein expression in stroke brains compared with controls (*p* = 0.0045 for the 57-kDa isoform and *p* < 0.0001 for both the 53-kDa and 34-kDa isoforms) (**Fig. 3D**).

**Fig. 3.**
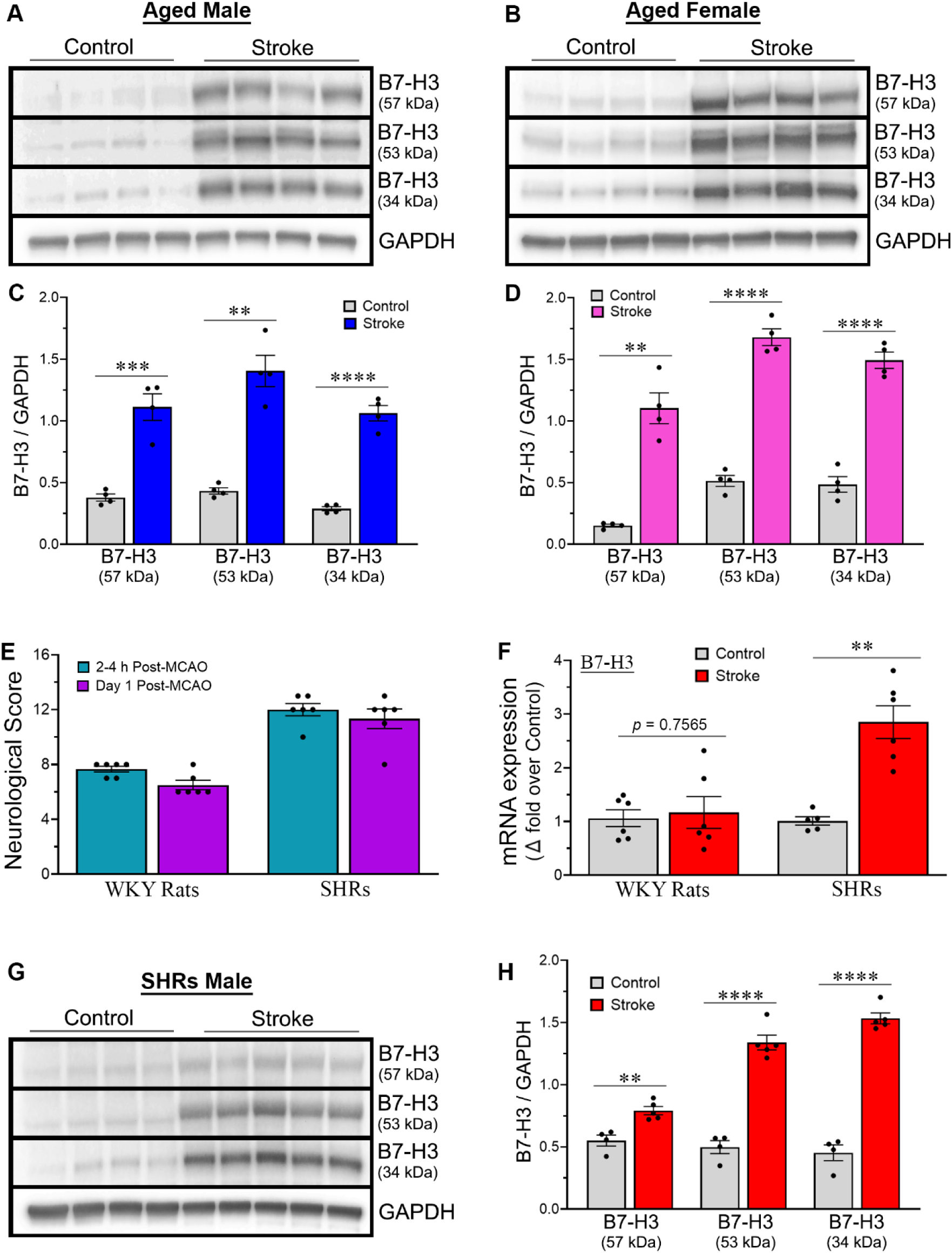
B7-H3 expression in the brain after ischemic stroke in clinically relevant rodent stroke models. (A, B) Immunoblots demonstrate upregulation of three B7-H3 isoforms (57, 53, and 34 kDa) in the ipsilateral (ischemic) brains of aged male (A) and female (B) mice compared with age-matched controls. Mice were subjected to 1-hour MCAO followed by reperfusion, and PBS-perfused brain tissue was collected on day 1 post-MCAO. (C, D) Column scatter plots show the quantified expression of the 57-, 53-, and 34-kDa B7-H3 isoforms relative to GAPDH expression in aged male (C) and female (D) mice. Error bars indicate SEM; n=4/group. \*\**p*<0.01, \*\*\**p*<0.001, \*\*\*\**p*<0.0001. (E) Column scatter plot showing modified neurological severity scores (mNSS) of male WKY rats and SHRs subjected to 1-hour MCAO followed by reperfusion. Error bars indicate SEM; n=6/group. (F) Column scatter plot shows B7-H3 mRNA expression on post-ischemic day 3 in male WKY rats and SHRs. Rats were subjected to 1-hour MCAO followed by reperfusion or sham surgery, and PBS-perfused ipsilateral brain tissue was collected on day 3. Error bars indicate SEM; n=5–6/group. \*\**p*<0.01. (G) Immunoblots demonstrate upregulation of three B7-H3 isoforms (57, 53, and 34 kDa) in the ischemic brains of male SHRs compared with controls. SHRs were subjected to 1-hour MCAO followed by reperfusion, and PBS-perfused brain tissue was collected on day 3 post-MCAO. (H) Column scatter plot shows the quantified expression of the 57-, 53-, and 34-kDa B7-H3 isoforms relative to GAPDH expression in SHRs. Error bars indicate SEM; n=4-5/group. \*\**p*<0.01, \*\*\*\**p*<0.0001.

Normotensive Wistar-Kyoto (WKY) rats are the standard control strain for spontaneously hypertensive rats (SHRs). Following 1-hour MCAO, modified neurological severity scores (mNSS) were 7.67 ± 0.21 and 6.5 ± 0.34 at 2-4 hours and day 1 post-MCAO, respectively, in WKY rats (**Fig. 3E**). In contrast, SHRs exhibited markedly higher mNSS scores of 12.0 ± 0.45 and 11.33 ± 0.71 at 2-4 hours and day 1 post-MCAO, respectively, indicating that an identical duration of MCAO produced more severe neurological deficits in hypertensive rats. A 1-hour MCAO in young male WKY rats and SHRs led to increased B7-H3 mRNA expression in the ipsilateral brain on day 3 post-MCAO in SHRs, but not in WKY rats (**Fig. 3F**). A two-tailed unpaired t-test (with Welch’s correction applied where appropriate) confirmed that B7-H3 mRNA levels were significantly higher in stroke-induced SHRs than in their control counterparts on day 3 (*p* = 0.0013). Western blot analysis of ipsilateral brain samples collected on day 3 post-MCAO from male SHRs showed increased protein expression of all B7-H3 isoforms in the stroke group compared with the control group (**Fig. 3G**). Two-tailed unpaired t-tests revealed that B7-H3 protein levels were significantly elevated in stroke-induced SHRs (*p* = 0.0029 for the 57-kDa isoform and *p* < 0.0001 for both the 53-kDa and 34-kDa isoforms) (**Fig. 3H**).

### 3.4. Effect of species, sex, age, and time after stroke on B7-H3 degradation in the ischemic brain

The ratio of expression of the 57-kDa glycosylated full-length membrane-bound 2IgB7-H3 isoform to that of the 34-kDa soluble B7-H3 (sB7-H3) isoform, which is potentially generated through proteolytic cleavage of the 57-kDa isoform, was calculated in the ipsilateral (ischemic) brains of rats and mice subjected to cerebral I/R. An increase in this 57/34-kDa ratio suggests reduced proteolytic cleavage or degradation of B7-H3 whereas a decrease in the ratio suggests enhanced degradation. Two-tailed unpaired t-test (with Welch’s correction applied when variances were unequal) showed that the 57/34-kDa ratio was significantly higher in young male mice (*p* = 0.0221) compared with age- and sex-matched SD rats on day 3 (**Fig. 4A**); in young female mice (*p* = 0.0038) compared with age-matched male mice on day 1 (**Fig. 4B**); in aged male mice (*p* = 0.0034) compared with sex-matched young mice on day 1 (**Fig. 4C**); and in young male mice on day 3 (*p* = 0.0058) compared with age- and sex-matched mice on day 1 (**Fig. 4D**). In contrast, the 57/34-kDa ratio was significantly lower in aged female mice (*p* = 0.0356) compared with age-matched male mice on day 1 (**Fig. 4B**) and in aged female mice (*p* = 0.0127) compared with sex-matched young mice on day 1 (**Fig. 4C**). Thus, the findings of this study demonstrate that species, sex, age, and time after stroke influence the extent of B7-H3 degradation in the ischemic brain following cerebral I/R.

**Fig. 4.**
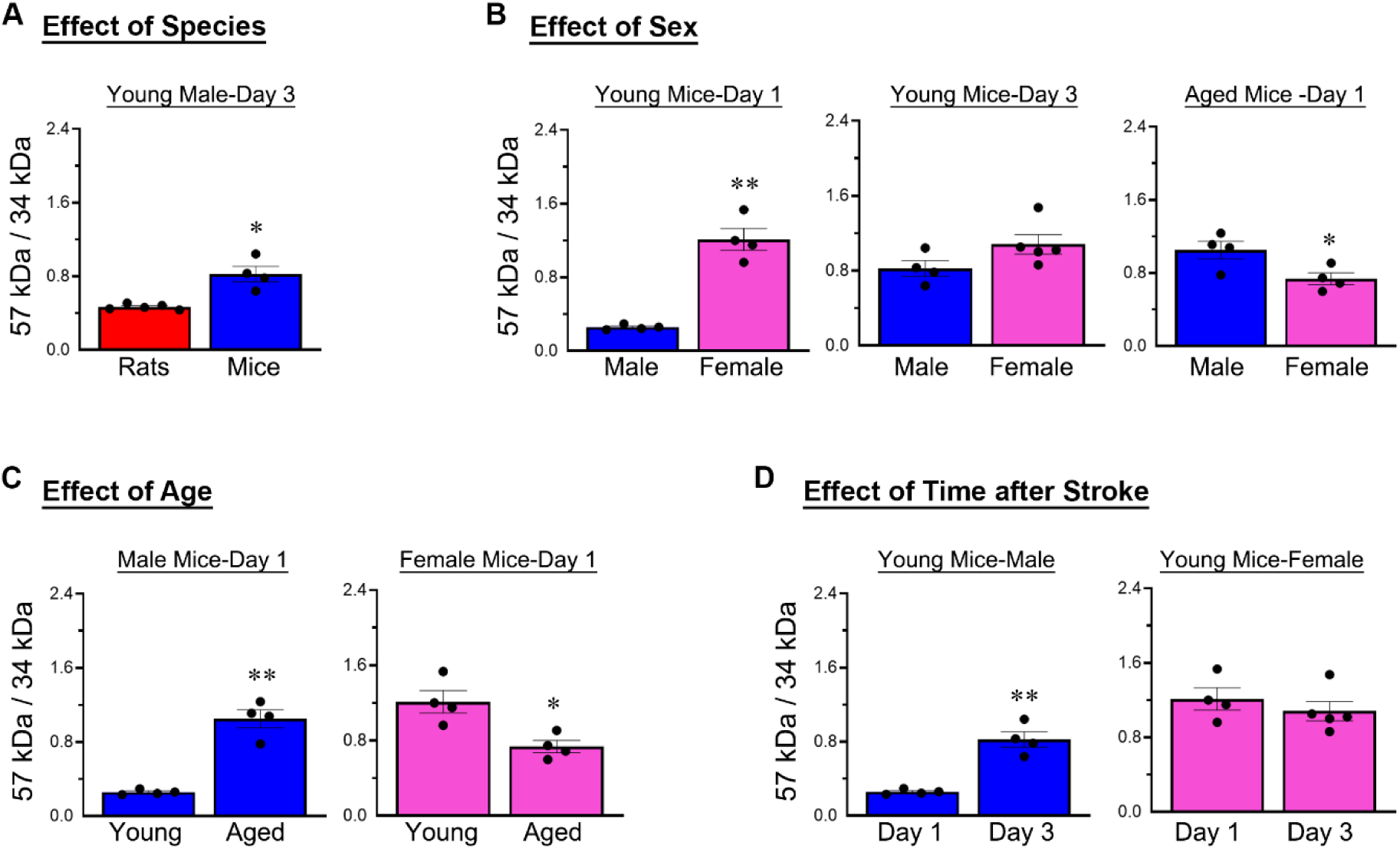
Species, sex, age, and time after stroke influence the extent of B7-H3 degradation in the ischemic brain after cerebral I/R. Column scatter plots illustrate the effects of species (A), sex (B), age (C), and time after stroke (D) on the ratio of expression of the 57-kDa to the 34-kDa B7-H3 isoform in the ipsilateral (ischemic) brain. Error bars indicate SEM; n=4-5/group. \**p*<0.05, \*\**p*<0.01 versus the corresponding experimental group.

### 3.5. Increased B7-H3 immunoreactivity is largely restricted to the ipsilateral ischemic hemisphere

Immunohistochemical analysis of coronal brain sections from SD rats collected on day 3 post-MCAO revealed prominent B7-H3 immunoreactivity in the brain (**Fig. 5A**). In control animals, B7-H3 staining was minimal in both hemispheres. In contrast, stroke animals exhibited robust B7-H3 staining that was predominantly confined to the ipsilateral (ischemic) hemisphere, particularly in the somatosensory cortex and caudate putamen (striatum), the primary regions affected by MCAO. Quantitative analysis showed that the B7-H3-positive area comprised approximately 23% of the ipsilateral hemisphere in the stroke group versus 0.1% in the control group (**Fig. 5B**). These findings suggest that B7-H3 upregulation is specific to ischemic injury and not attributable to surgical manipulation or procedural stress. Two-way ANOVA with Sidak’s multiple comparisons test showed that B7-H3 immunoreactivity, quantified as integrated density, was significantly greater in the ipsilateral hemisphere than in the contralateral hemisphere within the stroke group, and was also significantly greater in the ipsilateral hemisphere of stroke animals compared with the ipsilateral hemisphere of control animals (*p* < 0.0001 for both comparisons).

**Fig. 5.**
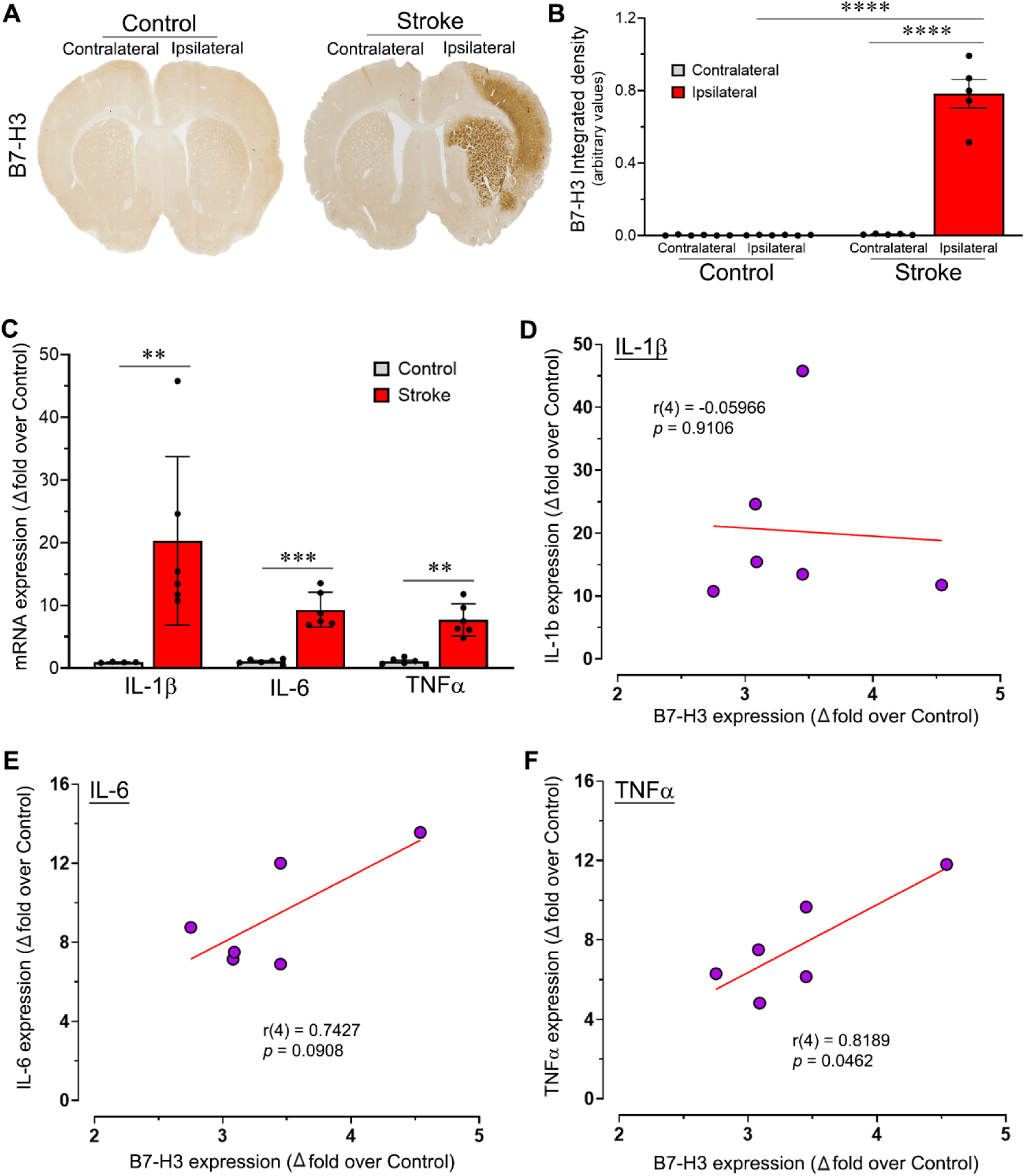
Post-stroke upregulation of B7-H3 and pro-inflammatory cytokines in the brain and correlation between B7-H3 and cytokine expression in the ischemic brain. (A) Representative coronal brain sections from young male Sprague-Dawley rats transcardially perfused with PBS followed by 10% neutral buffered formalin and collected on day 3 post-MCAO. Sections illustrate B7-H3 staining with DAB in control animals and in rats subjected to 2-hour MCAO followed by reperfusion. B7-H3 immunoreactivity is predominantly localized to the ipsilateral (ischemic) hemisphere, particularly in the somatosensory cortex and striatum, which are primary regions affected by MCAO. (B) Column scatter plot showing the quantified integrated density of B7-H3 staining in the contralateral (non-ischemic) and ipsilateral (ischemic) hemispheres of control and stroke groups. Error bars indicate SEM; n=5-6/group. \*\*\*\**p*<0.0001. (C) Column scatter plot showing mRNA expression of the pro-inflammatory cytokines IL-1β, IL-6, and TNFα in the ipsilateral brains of rats on day 3 post-MCAO. Error bars indicate SEM; n=4–6/group. \*\**p*<0.01; \*\*\**p*<0.001. (D-F) Scatter plots show the relationship between B7-H3 mRNA expression and the mRNA expression of IL-1β (D), IL-6 (E), and TNFα (F) in the ipsilateral brain on day 3 post-MCAO in stroke-induced rats.

### 3.6. B7-H3 expression exhibits a positive correlation with TNFα expression

Cerebral I/R induced by MCAO in young male SD rats increased the mRNA expression of the pro-inflammatory cytokines IL-1β, IL-6, and TNFα in the ipsilateral brain on day 3 post-MCAO (**Fig. 5C**). Statistical analysis showed that cytokine expression was significantly elevated in the stroke group compared with the control group for IL-1β (*p* = 0.0095; Mann-Whitney test), IL-6 (*p* = 0.0007; two-tailed unpaired t-test with Welch’s correction), and TNFα (*p* = 0.0013; two-tailed unpaired t-test with Welch’s correction). We next examined the relationship between B7-H3 mRNA expression and the mRNA expression of these pro-inflammatory cytokines on day 3 post-MCAO in stroke-induced SD rats. B7-H3 expression showed positive correlations with IL-6 and TNFα, but not with IL-1β (**Fig. 5D, E, F**). However, the correlation was statistically significant only between B7-H3 and TNFα expression (Pearson’s r(4) = 0.8189, *p* = 0.0462), indicating that higher B7-H3 expression is associated with higher TNFα expression in the ischemic brain.

## 4. Discussion

To our knowledge, this study is the first to demonstrate that B7-H3 expression is upregulated in the ischemic brain after cerebral I/R in two rodent species, encompassing males and females, young and aged animals, and animals with hypertension, a prevalent comorbidity among stroke patients. B7-H3 upregulation was largely confined to the ipsilateral (ischemic) hemisphere and was positively correlated with the expression of the pro-inflammatory cytokine TNFα. In young male SD rats, B7-H3 mRNA levels exhibited a time-dependent increase over the first week after ischemic stroke, peaking on day 3 and paralleling the time window associated with secondary ischemic injury. Moreover, B7-H3 protein levels were significantly elevated in the ipsilateral hemisphere at this time point in rats coinciding with the peak of mRNA expression. Although the temporal mRNA expression profile of B7-H3 in young male SD rats showed elevated B7-H3 mRNA levels exclusively on days 3 and 5 post-cerebral I/R, upregulation of B7-H3 isoform protein expression was evident as early as day 1 post-cerebral I/R in both sexes of young and aged mice. This observation may reflect the more pronounced stroke severity induced by the 1-hour MCAO in mice relative to the 2-hour MCAO in SD rats.

Immune checkpoints regulate the onset, severity, and duration of immune responses by mediating activating and inhibitory signals (Carlino et al., 2021). Among the 10 members of the B7 superfamily of immune checkpoints, B7-H3, a 316-amino-acid type 1 transmembrane protein, has attracted considerable interest since its discovery in 2001 (Chapoval et al., 2001; Liu et al., 2021). Although soluble B7-H3 is known to bind CD4+ T cells, CD8+ T cells, natural killer (NK) cells, and natural killer (NKT) cells, the specific receptor for B7-H3 remains unidentified (Kontos et al., 2021). Due to alternative splicing and post-translational modifications (PTMs) such as glycosylation, B7-H3 exists in multiple isoforms with different molecular weights. In humans, the main isoforms of B7-H3 include the full-length membrane-associated 4Ig-B7-H3 (∼45-66 kDa), the alternatively spliced 2Ig-B7-H3 variant (∼34-57 kDa), and a soluble form, sB7-H3 (∼37 kDa). While the 2Ig-B7-H3 isoform contains a single pair of extracellular V- and C-like Ig domains, a transmembrane region, and a cytoplasmic tail, the dominantly expressed human 4Ig-B7-H3 isoform contains two identical pairs of V- and C-like Ig domains (Chapoval et al., 2001; Sun et al., 2002). Previous work has shown that the 4Ig-B7-H3 isoform suppresses cytokine expression and inhibits T-cell proliferation, whereas the 2Ig-B7-H3 isoform enhances T-cell proliferation and upregulates IL-2 and IFNγ expression (Sun et al., 2011). Within the brain, cerebral microvascular endothelial cells may express low levels of B7-H3 mRNA; however, B7-H3 protein expression is minimal or undetectable in the normal brain (Digregorio et al., 2021). We hypothesize that brain cell types beyond cerebral microvascular endothelial cells may represent additional sources of B7-H3 in the brain following cerebral I/R. In this study, three predominant B7-H3 isoforms were detected by Western blot analysis in ipsilateral (ischemic) brain samples from both rats and mice subjected to cerebral I/R. A mouse monoclonal B7-H3 antibody raised against amino acids 166-465 within the N-terminal extracellular domain of human B7-H3 was used for detection. The observed B7-H3 isoforms had approximate molecular weights of 57, 53, and 34 kDa. Nevertheless, it has been reported that murine B7-H3 exists as a single isoform (2Ig-B7-H3), whereas human B7-H3 comprises two isoforms (2Ig-B7-H3 and 4Ig-B7-H3) (Ling et al., 2003; Sun et al., 2011). The three predominant isoforms detected at 57, 53, and 34 kDa in the ischemic brains of rats and mice may arise from alternative splicing of B7-H3 mRNA and/or PTMs of the B7-H3 protein. PTMs enhance the functional diversity of proteins, can affect normal cellular function, and may contribute to disease pathogenesis (Getu et al., 2023). The best-characterized PTM of B7-H3 is glycosylation. During glycosylation, carbohydrate chains are covalently attached to the asparagine motifs of B7-H3, leading to an increase in molecular weight. Thus, the molecular weights of the glycosylated 4Ig-B7-H3 and 2Ig-B7-H3 isoforms in humans are approximately 90-100 kDa and 50-70 kDa, respectively. In mice, the calculated molecular weight of the full-length 2Ig-B7-H3 isoform is approximately 25-26 kDa. The isoform observed at 57-kDa in this study within the ipsilateral (ischemic) brains of rodents may correspond to the N-glycosylated full-length membrane-associated 2Ig-B7-H3 isoform. The isoforms detected at 53 kDa and 34 kDa could have been generated through either alternative splicing or proteolytic cleavage of the N-glycosylated full-length, membrane-bound 57 kDa 2Ig-B7-H3 isoform. The isoform observed at 34 kDa may correspond to the soluble form of B7-H3 (sB7-H3), potentially arising from either proteolytic cleavage of the glycosylated full-length, membrane-associated 2Ig-B7-H3 isoform or alternative splicing of B7-H3 mRNA (Sun et al., 2011). Although the production of sB7-H3 via alternative mRNA splicing remains unclear, cleavage of the extracellular domain of the membrane-bound glycosylated 2Ig-B7-H3 by matrix metalloproteinases (MMPs) has been recognized as the primary mechanism underlying the formation of sB7-H3 (Zhang et al., 2008). If sB7-H3 levels increase in serum after ischemic stroke, it could serve as a clinically valuable biomarker for assessing stroke severity, neuroinflammatory burden, or therapeutic response.

The ratio of the 57-kDa to the 34-kDa B7-H3 isoforms, which primarily reflects the proteolytic cleavage/degradation status of B7-H3, demonstrated that species, sex, age, and time post-stroke influence the degree of B7-H3 degradation in the ischemic brain following cerebral I/R. As described earlier, an increase in 57/34-kDa indicates reduced proteolytic cleavage or degradation of B7-H3, and a decreased ratio indicates enhanced degradation. Across experimental groups, B7-H3 degradation was lower in mice than in rats. Among young mice on post-ischemic day 1, females exhibited less degradation (higher 57/34-kDa ratios) than males. In contrast, among aged mice on day 1, females showed greater degradation than males. When age groups were compared within sex on day 1, aged males displayed less degradation than young males, whereas aged females showed greater degradation than young females. Finally, in young male mice, B7-H3 degradation on post-ischemic on day 3 was lower than in age- and sex-matched mice on day 1. These variations in B7-H3 degradation likely reflect a combination of species and sex differences, age-related factors, differences in stroke severity, and time-dependent changes in the expression of various MMPs following cerebral I/R. Our previous work demonstrating distinct temporal patterns of MMP expression in the post-ischemic brain, together with the present findings, support the notion that MMPs contribute to the proteolytic cleavage of membrane-bound B7-H3 into its soluble form (Zhang et al., 2008; Chelluboina et al., 2015). Although not directly tested here, it will be important in future studies to determine whether B7-H3 isoform levels or 57/34-kDa degradation ratios correlate with stroke severity scores (e.g., mNSS) across different experimental groups.

Hypertension is a significant risk factor for ischemic stroke and, as a comorbid condition, increases both the incidence and severity of stroke (Fisher et al., 2007; Howells et al., 2010; O’Donnell et al., 2010). The SHR is the most widely used genetic model of hypertension. In comparison with normotensive wild-type WKY rats, SHRs exhibit consistently elevated blood pressure beginning at approximately 4 weeks of age (Dickhout and Lee, 1998). In this study, 3-month-old SHRs were subjected to cerebral I/R. A 1-hour MCAO produced more severe neurological deficits in SHRs than in normotensive WKY rats as reflected by mNSS values, illustrating the impact of hypertension on the severity of brain damage following cerebral I/R. Furthermore, stroke severity, as measured by mNSS, was greater after 1-hour MCAO in SHRs than after 2-hour MCAO in normotensive SD rats. As expected, B7-H3 mRNA expression was significantly increased in SHRs compared with wild-type WKY rats. In addition, protein expression levels of all B7-H3 isoforms were markedly elevated in stroke-induced SHRs relative to control SHRs. Consistent with our findings in SD rats, the expression level of the 57-kDa B7-H3 isoform in SHRs on day 3 following cerebral I/R was lower than the expression levels of the 53- and 34-kDa isoforms.

The robust B7-H3 immunoreactivity observed in the cortex and striatum of the ipsilateral hemisphere, the primary regions impacted by MCAO, together with undetectable or very low levels of B7-H3 immunoreactivity in the contralateral hemisphere, suggests a potential role for B7-H3 in stroke pathology. The increased B7-H3 expression in the ipsilateral brain following cerebral I/R was positively correlated with the expression of the key pro-inflammatory cytokine TNFα, suggesting that B7-H3 may participate in the inflammatory cascade after cerebral I/R. This finding is consistent with a prior study indicating that B7-H3 can promote the release of pro-inflammatory cytokines from monocytes and macrophages (Zhang et al., 2010). Thus, our data provide evidence that B7-H3 is not only upregulated in the ischemic brain but may also be linked to pro-inflammatory cytokine signaling. However, these observations do not establish whether elevated B7-H3 expression in the ischemic brain is ultimately harmful or beneficial. The correlation between B7-H3 and TNFα after cerebral I/R may also indicate that B7-H3 expression increases to protect brain cells from excessive neuroinflammation. For example, in the tumor microenvironment, B7-H3 promotes macrophage polarization from an M1 (pro-inflammatory) to an M2 (anti-inflammatory) phenotype, contributing to an immunosuppressive microenvironment through the CCL2-CCR2-M2 macrophage axis. B7-H3 further enhances immunosuppression by increasing IL-10 and TGFβ production, inhibiting the activity of multiple T cell subsets and other immune cells, and reducing the secretion of IFNγ, IL-2, perforin, and granzyme B (Mao et al., 2017; Han et al., 2018; Lu et al., 2020; Lee et al., 2021; Long et al., 2021; Si et al., 2022; Miyamoto et al., 2022). Although B7-H3 is recognized as an immunoregulatory molecule with context-dependent co-stimulatory or co-inhibitory functions, its role in neuroinflammation after cerebral I/R remains unknown. In the ischemic microenvironment, B7-H3 may therefore act either as a friend or as a foe.

In summary, B7-H3 expression is significantly upregulated in the ischemic brains of young rats and of both young and aged male and female mice. B7-H3 immunoreactivity is largely confined to the ipsilateral (ischemic) hemisphere and is particularly prominent in the cortex and striatum, the primary regions affected by MCAO, suggesting a role in stroke pathology. Although B7-H3 is best known for its role in immune regulation, its increased expression following stroke highlights its potential as a therapeutic target. The specific role of B7-H3 in neuroinflammation and brain damage after cerebral I/R, however, remains unknown. Elucidating the function of B7-H3 in the ischemic brain microenvironment may lead to the discovery of novel targets and the development of treatments that reduce brain damage and enhance functional recovery following cerebral I/R. In this study, B7-H3 expression in the ischemic brains of rats was positively correlated with expression of the pro-inflammatory cytokine TNFα, raising the possibility that blocking B7-H3 could mitigate post-ischemic neuroinflammation, a key contributor to secondary brain injury. B7-H3 may also serve as a marker of neuroinflammatory burden in the brain after cerebral I/R. The scope of this study was limited to demonstrating B7-H3 upregulation in the brain following cerebral I/R in preclinical rodent models. Future studies will focus on identifying the cellular sources of B7-H3 in the ischemic brain and characterizing its role in post-stroke pathogenesis, particularly neuroinflammation, brain damage, and the recovery of sensorimotor and cognitive functions. These studies will use genetic and pharmacological approaches in aged animals and in animals with comorbidities to better reflect the heterogenous health status of the acute ischemic stroke patient population.

## Author contributions

Conceptualization, funding acquisition, and supervision: K.K.V., S.A., J.D.K.; Animal surgeries and care: S.R.C., I.M.B., and K.K.V.; Methodology, investigation, and data acquisition: S.R.C., I.M.B., C.A.F., S.R.M., N.K., and S.J.; Data analysis and interpretation: K.K.V., C.A.F., and S.R.C.; Manuscript preparation: K.K.V.; Manuscript editing: C.A.F.; Manuscript review and final approval: All authors; Both S.R.C. and I.M.B. contributed equally to this work and share first authorship.

## Acknowledgments

We thank the National Institute of Neurological Disorders and Stroke of the National Institutes of Health for financial support.

